# Two-dimensional multiplexed assay for rapid and deep SARS-CoV-2 serology profiling and for machine learning prediction of neutralization capacity

**DOI:** 10.1101/2021.08.03.454782

**Authors:** Akiko Koide, Tatyana Panchenko, Chan Wang, Sara A. Thannickal, Larizbeth A. Romero, Kai Wen Teng, Francesca-Zhoufan Li, Padma Akkappedi, Alexis D. Corrado, Jessica Caro, Catherine Diefenbach, Marie I. Samanovic, Mark J. Mulligan, Takamitsu Hattori, Kenneth A. Stapleford, Huilin Li, Shohei Koide

## Abstract

Antibody responses serve as the primary protection against SARS-CoV-2 infection through neutralization of viral entry into cells. We have developed a two-dimensional multiplex bead binding assay (2D-MBBA) that quantifies multiple antibody isotypes against multiple antigens from a single measurement. Here, we applied our assay to profile IgG, IgM and IgA levels against the spike antigen, its receptor-binding domain and natural and designed mutants. Machine learning algorithms trained on the 2D-MBBA data substantially improve the prediction of neutralization capacity against the authentic SARS-CoV-2 virus of serum samples of convalescent patients. The algorithms also helped identify a set of antibody isotype–antigen datasets that contributed to the prediction, which included those targeting regions outside the receptor-binding interface of the spike protein. We applied the assay to profile samples from vaccinated, immune-compromised patients, which revealed differences in the antibody profiles between convalescent and vaccinated samples. Our approach can rapidly provide deep antibody profiles and neutralization prediction from essentially a drop of blood without the need of BSL-3 access and provides insights into the nature of neutralizing antibodies. It may be further developed for evaluating neutralizing capacity for new variants and future pathogens.

## INTRODUCTION

COVID-19 remains a major threat to the entire world. Although multiple effective vaccines against SARS-CoV-2 have been deployed (Baden et al., 2021; Polack et al., 2020; Voysey et al., 2021), there remains intense interest in characterizing the anti-SARS-CoV-2 immunity in terms of humoral and cellular immune responses. A number of serology testing methods have been deployed for the purposes of determining binary seroconversion as the indication of past exposure to the virus and of analyzing antibody responses (Amanat et al., 2020; den Hartog et al., 2020; Dobano et al., 2021; Dogan et al., 2021; Garcia-Basteiro et al., 2020; Marien et al., 2021; Pisanic et al., 2020; Weiss et al., 2020; Yates et al., 2021). In parallel, a number of assays have been developed to determine the neutralization capacity against the virus in cellular entry, the primary protective role of humoral responses. Neutralization assays using authentic SARS-CoV-2 viruses, including plaque reduction neutralization test (PRNT) and microneutralization assay (MNA), are clearly the gold standard (Bewley et al., 2021), but they are cell-based assays that need to be performed in a BSL-3 environment and are inherently low-throughput. Many studies, including those cited above, have demonstrated good correlation between antibody levels and neutralization capacity as one would expect. However, the degree of correlation from a single measurement is still moderate. Few have reported attempts to improve the prediction of neutralization from antibody levels. The ability to accurately predict neutralization capacity from antibody profiles, which can be rapidly measured outside a BSL-3 environment, could enrich the value of antibody profiling and potentially lay a foundation for simpler assays, e.g., point-of-care tests, for predicting immunity against SARS-CoV-2.

SARS-CoV-2 encodes four structural proteins and additional nonstructural proteins after proteolytic processing (Bojkova et al., 2020), and SARS-CoV-2 infection elicits multiple antibody isotypes, i.e., IgM, IgA and IgG, binding to many of these viral proteins. Antibodies to the N-protein (NP) was used as the antigen in clinical tests for past infection. As expected from the localization of NP inside the viral particle, antibodies to NP do not neutralize viral entry and their levels are poorly correlated to neutralization titers (Noval et al., 2021).

Spike is the major surface antigen of SARS-CoV-2 that plays the central role in the attachment to and entry into cells (Hoffmann et al., 2020; Walls et al., 2020). Its receptor-binding domain (RBD) interacts with the canonical receptor, angiotensin-converting enzyme 2 (ACE2) (Hoffmann et al., 2020; Lan et al., 2020; Walls et al., 2020). Consequently, antibodies from infected subjects, as well as engineered reagents that interfere with the RBD-ACE2 interaction, can inhibit viral infection (Hansen et al., 2020; Hattori et al., 2021; Ju et al., 2020; Robbiani et al., 2020; Shi et al., 2020; Wec et al., 2020; Wu et al., 2020). Accordingly, most current vaccines utilize the prefusion-stabilized form of spike as the antigen (Baden et al., 2021; Polack et al., 2020).

Because of the importance of the RBD in viral infection, early studies focused on antibodies directed to the RBD. Indeed, most of potent neutralizing antibodies, including those that have been authorized for clinical use, bind to the RBD and interfere with the interaction of the RBD with ACE2. However, Spike is a large protein, and natural infection and vaccination elicit antibodies to diverse regions of Spike (Amanat et al., 2021; Voss et al., 2021). Intriguingly, there are neutralizing antibodies that bind to the spike protein outside the RBD, e.g. the N-terminal domain (Brouwer et al., 2020; Voss et al., 2021). Consequently, anti-RBD antibodies may not adequately account for all neutralizing antibodies in many cases.

A number of proxy assays, including those using pseudotyped viruses and engineered cell lines, have been developed that do not require the authentic SARS-CoV-2 virus or a BSL-3 environment (Nie et al., 2020; Riepler et al., 2020; Tan et al., 2020). Although they substantially increase the access and throughput of neutralization evaluation, they show only moderate concordance with the gold-standard assay (Noval et al., 2021), possibly because these systems do not fully recapitulate molecular interactions during viral entry. Furthermore, because serology assays and neutralization assays are based on distinct principles, there is a paucity of simple and rapid methods for quantifying *both* antibody profiles and neutralizing capacity of patient samples, although there is strong correlation between antibody levels and neutralization, as expected.

We envisioned that we can quantitatively predict the viral neutralization capacity by examining the antibody profile in patient samples more deeply and quantitatively in terms of antigen, epitope and antibody type. Thus, we have developed a multiplex assay for SARS-CoV-2 serology that enables us to deeply characterize the antibody profile in terms of antibody isotypes and epitopes, and applied machine learning to develop predictive algorithms of neutralization capacity.

## RESULTS

### Assay design and validation

To improve on our one-dimensional multiplex bead-based binding assay (MBBA) (Hattori et al., 2020), we have developed two-dimensional multiplex bead-based binding assays (2D-MBBA) combined with flow cytometry detection to profile antibody isotypes for multiple antigens that are site-specifically biotinylated (Fig. 1A and 1B). This new method can simultaneously detect a total of 15 (5 x 3) antibody-antigen interactions in a single reaction (Fig. 1B). We tested commercially available fluorescent microbeads (Qbeads) and also Dynabeads M-280 streptavidin that were fluorescently labeled in-house using a biotin-dye conjugates (Hattori et al., 2020). Both beads types allowed us to produce a total of five distinct levels of fluorescence intensity for multiplexing. We used commercially available secondary antibodies for human IgG, IgM and IgA that are labeled with different dyes. Although both platforms were capable of performing the multiplex assays, Qbeads showed significantly lower background binding to serum samples, and thus we chose Qbeads for subsequent experiments. Although it is common to use dye combinations that require compensation for detecting multiple antibodies, we found it difficult to accurately measure the levels of each antibody isotype using such dye combinations because many of serum/plasma samples have large signal for one isotype while very small signal for other isotypes. Therefore, for simultaneously and accurately detecting antibody isotypes, we chose a set of fluorescent dye-labeled secondary antibodies that requires no signal compensation on an appropriately equipped flow cytometer (see Methods).

**Figure 1.**
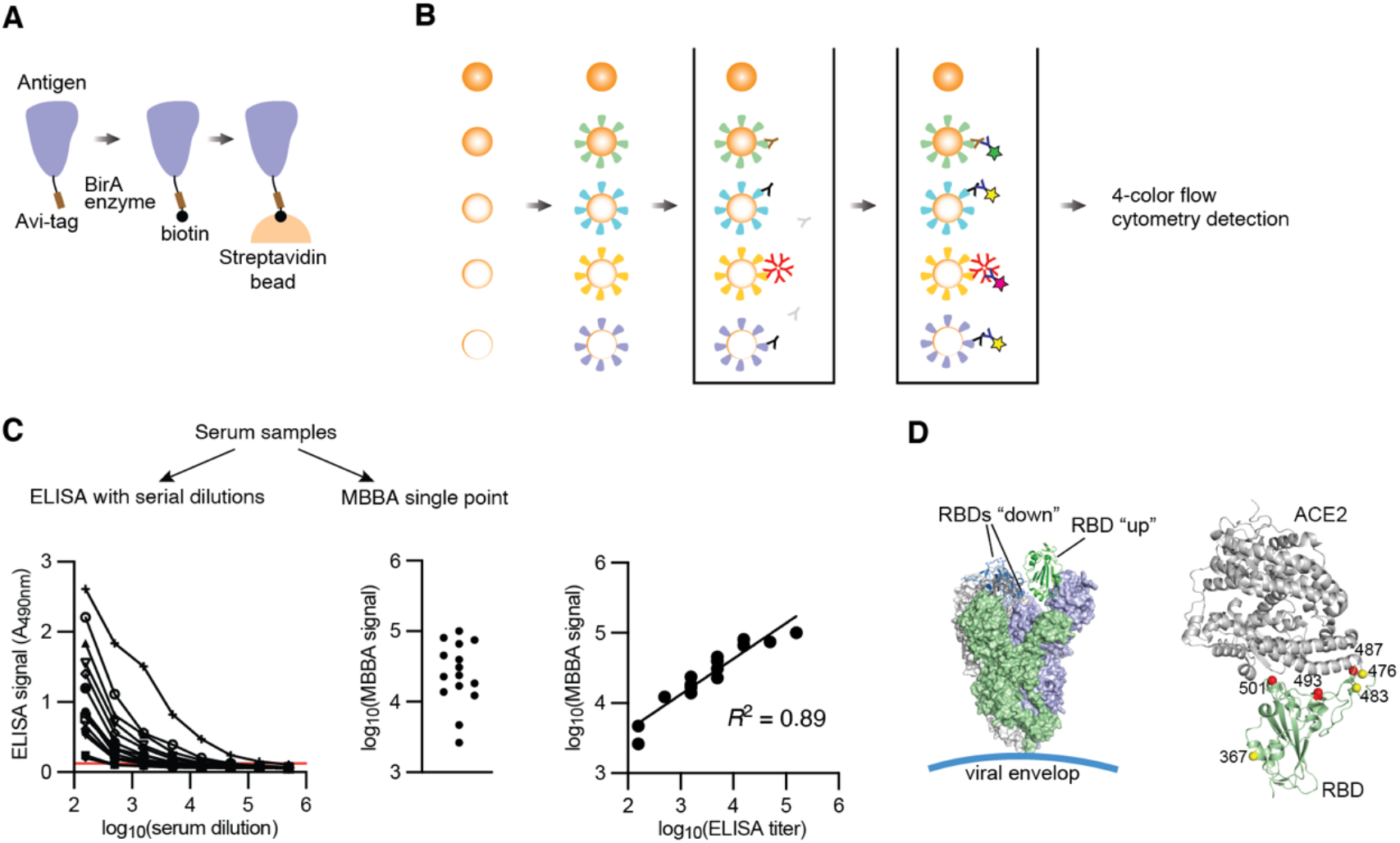
Design of 2D-MBBA for SARS-CoV-2 antibody profiling. (A) Scheme showing the design of an Avi-tagged antigen and site-specific immobilization on a streptavidin-coated bead. (B) Scheme showing the 2D-MBBA principle. Five types of microbeads each presenting a different antigen are mixed and reacted with a serum sample. After washing, bound antibodies are detected with isotype-specific secondary antibodies on a four-color flow cytometry that separately quantify the beads and three isotypes. (C) Comparison of MBBA with conventional ELISA using serum samples and RBD. ELISA end point titers were determined using the cutoff shown as the red horizontal line. The *R*^2^ values were determined after log10 transformation of the ELISA endpoint titers and MBBA signals. (D) Antigen design. Schematic drawings of the spike protein (PDB ID: 6VSB) and of RBD in complex with ACE2 (PDB ID: 6VW1) denoting mutations used in this study.

Compared with a standard ELISA, our multi-dimensional approach has a number of advantages: (i) viral antigens are presented on microspheres via a flexible linker, which minimizes masking of epitopes due to preferential binding to plastic surface (Fig. 1A); (ii) multiple antigens are pooled in a single reaction; (iii) multiple antibody isotypes (IgG, IgM and IgA) are simultaneously detected; (iv) signals are not dependent on an enzyme reaction and thus time-independent; and (v) the sensitivity and dynamic range of flow cytometry far exceed those of ELISA, eliminating the need for measuring serially diluted serum samples. Consequently, this method requires a few microliters of blood samples. These attributes in combination substantially improves the assay throughput using substantially less antigen proteins and blood samples compared with ELISA. Validation of this method is shown in Fig. S1.

Using a series of serum samples and the RBD as an antigen, we compared our method with the ELISA endpoint assay with serial dilutions (Fig. 1C), an approved standard assay for SARS-CoV-2 serology (Amanat et al., 2020). The single-point signals from our MBBA (i.e., no serial dilutions) showed excellent correlation (*R^2^* = 0.81) with the ELISA titer with eight dilutions, indicating that our method achieves a large dynamic range without the need to perform serial dilutions and reducing labor-intensive steps without compromising data quality. Because of the 15-plex capability (3 isotypes for 5 antigens), MBBA measurements using a 96-well plate is equivalent to 11,520 wells of single-plex ELISA wells.

To quantify antibodies binding to distinct regions of Spike, we designed and produced a series of SARS-CoV-2 antigens, including Spike, the RBD of Spike, and natural and designed RBD mutants (Fig. 1D, Supplementary Fig. 2). Because neutralizing antibodies should recognize the folded, prefusion conformation of Spike, we did not include short, unstructured fragments, although it is possible that some neutralizing antibodies may cross-react with such fragments. Natural RBD mutants included RBD-V483A, RBD-V367F and RBD-G476S that were in circulation in the U.S. in 2020, and a designed triple mutant that disrupts the RBD-ACE2 interaction, N487K, Q493K and N501K in Spike (termed Spike-T hereafter) and RBD (termed RBD-T hereafter; Fig. 1D) (Hattori et al., 2021). These designed mutations disrupt the interaction of RBD with neutralizing antibodies that target the ACE2-interaction surface. We also produced RBD of SARS-CoV1 (referred to as RBD-CoV1), which has 74.5 % identity with SARS-CoV-2 RBD. These antigens are all site-specifically biotinylated using the Avi-tag located in a C-terminal extension and immobilized on the beads via the strong interaction between biotin and streptavidin. The presentation of full-length spike using a C-terminal biotin coupled to streptavidin-coated beads approximate its orientation on the viral surface (Fig. 1A and 1D).

We next applied 2D-MBBA to measure antibody profiles of samples from 101 convalescent hospital workers and 34 negative control samples, the same sample set that we previously characterized using RBD and NP ELISA as well as a neutralization assay with the authentic virus (Noval et al., 2021). We characterized the binding of IgG, IgM and IgA to a total of eight antigens plus a no-antigen control (Fig. 2). Binding signals of the same antibody isotype (IgG, IgM or IgA) to SARS-CoV-2 antigens, i.e., Spike, RBD and RBD mutants, were well correlated (Fig. 2A), as one might expect from the fact that these antigens are all derived from the spike protein. In contrast, binding signals of different isotypes to a given antigen were not highly correlated, confirming diverse isotype profiles among these patients (Noval et al., 2021). Consistent with other studies (Long et al., 2020; Okba et al., 2020), these results establish that antibody profiling provides sensitive and specific diagnosis for seroconversion after SARS-CoV-2 infection (Fig. 2A). Specificities are over 0.95 for all the samples. Sensitivities for IgG for all the RBD variants of SARS-CoV-2 antigens and IgM for Spike are extremely high with values of 1.00 and 0.99, respectively.

**Figure 2.**
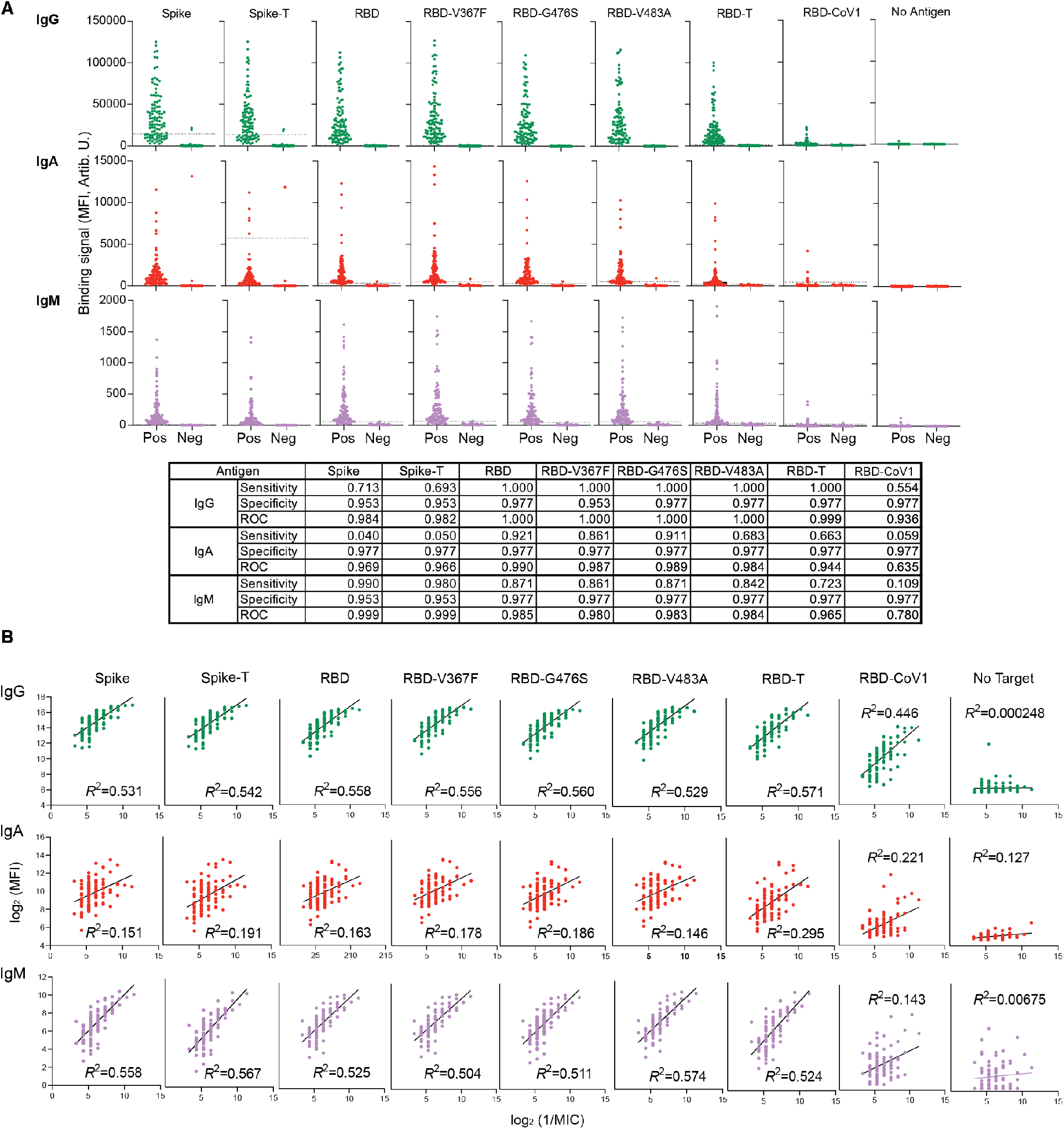
2D-MBBA data for convalescent (positive) and pre-COVID (negative) samples and the diagnostic performance of individual datasets. (A) Raw 2D-MBBA readouts of IgG, IgA and IgM levels for the indicated antigens of convalescent (“Pos”) and pre-COVID (“Neg”) samples. For each isotype-antigen pair, sensitivity and specificity for detecting positivity were calculated with cut-off values set at the mean + 3 x s.d. of the pre-COVID data. The ROC values denote the area under the curve for the ROC curves (ROC curves are included in Supplementary Data Table 1). (B) 2D-MBBA data for the convalescent sera plotted versus neutralization titer after log2 transformation. Note that the negative control data are not included, in order to critically evaluate the correlation of each isotype-antigen pair with neutralization.

We examined the correlation of individual isotype–antigen datasets with neutralization titers of these samples that we previously reported (Noval et al., 2021) (Fig. 2B). We intentionally excluded negative control samples in this analysis, in order to more critically evaluate the predictive power of these datasets. Still, most of individual datasets exhibit significant positive correlation with neutralization MIC. *R^2^* values for the IgG and IgM datasets analyzed with the simple linear regression model after logarithmic transformation were above 0.5. Although the signal strengths to RBD-CoV1 were substantially lower than those to RBD (Fig. 2A), the IgG–RBD-CoV1 dataset showed good correlation (*R*^2^ = 0.446) (Fig. 2B), which may reflect weak cross-reactivity of anti-RBD IgG to RBD-CoV1.

Interestingly, the correlations between neutralization titers and the IgG and IgM datasets to Spike-T and RBD-T, the designed mutant antigens harboring the triple mutants that disrupt the RBD-ACE2 interaction, were among the highest of all isotype-antigen pairs. These results are consistent with our previous results that revealed that the ACE2-interacting surface is not highly immunogenic (Hattori et al., 2021) and with other reports showing that a small fraction of antibodies target this surface of the spike protein (Voss et al., 2021).

The correlations of IgG and IgM datasets from 2D-MBBA and from our previous ELISA-based analysis (Noval et al., 2021) are consistent. In contrast, the IgA data from 2D-MBBA showed substantially weaker correlation than the ELISA data. We speculate that the MBBA format with low antibody density and more stringent washing removed a subset of IgA with low affinity, whereas ELISA with high antigen density tends not to discriminate low and high-affinity antibodies. IgA in the blood mostly does not have the J chain and thus it has an IgG-like bivalent architecture, unlike the tetravalent IgA in the mucosal tissues (Woof and Russell, 2011), and thus low-affinity IgA species in serum may be particularly sensitive to stringent binding conditions. One may be able to exploit this sensitivity to quantify high-affinity and low-affinity antibody species in future applications.

### Developing machine learning algorithms to predict neutralization capacity from serology data

We then tested whether we can more accurately predict neutralization by utilizing the entire 2D-MBBA datasets with machine learning algorithms. To this end, we subjected the dataset, consisting of 27 assays as input, to multiple well-known machine learning algorithms: stepwise linear regression (Miller, 2002), elastic net regression (Zou and Hastie, 2005), multivariate adaptive regression splines (MARS) (Friedman, 1991), classification and regression tree (CART) (Lewis, 2000), and Bayesian regularized neural networks (BRANNs) (Burden and Winkler, 2008). The predictive performance of these algorithms was evaluated by three metrics: root mean square error (RMSE), *R*^2^, and mean absolute error (MAE) with 10-fold crossvalidation (CV). The elastic net regression and BRANNs methods had equivalently best results with the smallest RMSE and MAE and the largest *R*^2^ (=0.73) (Table 1), substantially improving the prediction accuracy when they are compared with the predictions based on a single assay (Fig. 2).

**Table 1.** Performance of different machine learning algorithms in predicting the neutralization titer from 2D-MBBA data. The mean and standard deviation (in the parentheses) of the considered three metrics based on 100 independent replicates are reported.

| All data (27 pairs) |  | Method |  |  |  |  |
| --- | --- | --- | --- | --- | --- | --- |
| Group | Metric | Stepwise | Elastic net | MARS | CART | BRANNs |
| Test* | RMSE | 0.86 ( 0.21 ) | 0.81 ( 0.19 ) | 0.92 ( 0.23 ) | 1.07 ( 0.3 ) | 0.8 ( 0.2 ) |
| | $R^2$ | 0.69 ( 0.15 ) | 0.73 ( 0.14 ) | 0.65 ( 0.18 ) | 0.56 ( 0.2 ) | 0.73 ( 0.15 ) |
|  | MAE | 0.69 ( 0.18 ) | 0.66 ( 0.16 ) | 0.74 ( 0.19 ) | 0.87 ( 0.24 ) | 0.66 ( 0.17 ) |
| Training<sup>#</sup> | RMSE | 0.82 ( 0.04 ) | 0.8 ( 0.03 ) | 0.89 ( 0.04 ) | 1.03 ( 0.05 ) | 0.79 ( 0.02 ) |
| | $R^2$ | 0.73 ( 0.02 ) | 0.75 ( 0.02 ) | 0.67 ( 0.04 ) | 0.59 ( 0.04 ) | 0.75 ( 0.02 ) |
|  | MAE | 0.67 ( 0.03 ) | 0.66 ( 0.02 ) | 0.72 ( 0.03 ) | 0.84 ( 0.04 ) | 0.65 ( 0.02 ) |
| Reduced data (15 pairs) |  | Method |  |  |  |  |
| Group | Metric | Stepwise | Elastic net | MARS | CART | BRANNs |
| Test | RMSE | 0.85 ( 0.19 ) | 0.88 ( 0.19 ) | 0.9 ( 0.23 ) | 1.02 ( 0.25 ) | 0.8 ( 0.16 ) |
| | $R^2$ | 0.72 ( 0.16 ) | 0.72 ( 0.18 ) | 0.64 ( 0.22 ) | 0.57 ( 0.21 ) | 0.73 ( 0.17 ) |
|  | MAE | 0.69 ( 0.17 ) | 0.73 ( 0.16 ) | 0.72 ( 0.17 ) | 0.83 ( 0.2 ) | 0.66 ( 0.14 ) |
| Training | RMSE | 0.83 ( 0.03 ) | 0.82 ( 0.02 ) | 0.87 ( 0.04 ) | 1.02 ( 0.06 ) | 0.78 ( 0.02 ) |
| | $R^2$ | 0.73 ( 0.02 ) | 0.75 ( 0.02 ) | 0.69 ( 0.03 ) | 0.61 ( 0.04 ) | 0.76 ( 0.02 ) |
|  | MAE | 0.67 ( 0.03 ) | 0.67 ( 0.02 ) | 0.7 ( 0.03 ) | 0.82 ( 0.05 ) | 0.64 ( 0.02 ) |

\*We first trained the model using the training set, then calculated the predictive metrics with the trained model using the test set
<sup>#</sup>We trained the model and calculated the metrics based on the trained model using the same training set.

Next, we determined the importance of the 27 MBBA datasets by the regression coefficients in the elastic net regression and connection weights (Olden et al., 2004) in the BRANNs predictive models respectively. Because both regression coefficients and connection weights have the magnitude of importance (the absolute value) and also the direction of correction (the sign; positive or negative), we rank the importance by their absolute values. For BRANN, IgG–Spike-T, IgM–Spike-T and IgM–RBD-V483A were ranked as the top three (Fig. S4B). For Elastic Net, IgG–RBD-T, IgM–Spike-T and IgM–RBD-V483A were the top three. The selection of both RBD-T and Spike-T, which have mutations to eliminate antibody binding to the ACE2-interaction surface of the RBD (Hattori et al., 2021), may be rationalized based on our previous observation that a small fraction of RBD-binding antibodies target the ACE2-interaction surface, as discussed above, and by the notion that IgM binding to this surface should be captured in the data with RBD-V483A.

We then tested the effectiveness of algorithms using a reduced set of input data with the goal of establishing an assay that can be performed in a single well per sample (i.e., ≤5 antigens; Fig. 1A). We chose four antigens that included isotype–antigen datasets that are least correlated (Fig. S4A) and scored high in the importance analysis (Fig. S4B): Spike-T, RBD, RBD-V483A, RBD-T, plus a no-antigen control. We chose Spike-T over Spike from a practical reason. Although datasets for these two antigens are nearly perfectly correlated (Fig. S4A), the purified Spike-T protein was less prone to aggregation and had higher production yield than Spike, and thus the use of Spike-T may lead to more robust assay.

For the reduced datasets, we evaluated the predictive performance of the six machine learning algorithms in the same manner as for the full datasets. The reduction of the input datasets did not affect the prediction performance. BRANNs performed the best in terms of all three metrics (Table 1). These results confirm the redundancy of the input datasets and the ability of machine learning algorithms to achieve accurate prediction from a smaller number of input MBBA datasets.

### Testing the trained algorithms with data from vaccinated subjects

We next tested how well the algorithms trained with the convalescent samples predict the neutralization capacity of plasma samples from vaccinated cancer patients. Healthy people usually mount strong immune responses to the mRNA vaccines (Jackson et al., 2020; Walsh et al., 2020) and therefore neutralization prediction may have limited values. However, we and others have found that immune compromised patients tend to have limited immune responses to the vaccines (Agha et al., 2021; Boyarsky et al., 2021; Deepak et al., 2021; Diefenbach et al., 2021; Herishanu et al., 2021), and therefore accurate assessment of their antibody profiles and neutralization capacity may be particularly important for the protection of this vulnerable population.

Since the evaluation of the convalescent samples in mid 2020, we have modified our neutralization assay. Instead of a cell death-based assay, we used a fluorescent reporter protein-based assay for new samples (Muruato et al., 2020; Xie et al., 2020). We used a subset of the convalescent samples and confirmed that two assays are highly correlated and comparable (Supplementary Figure 2E).

We analyzed a total of 23 samples from vaccinated lymphoma patients (Diefenbach et al., 2021) with the reduced set antigens. The IgG–RBD and IgG–mutant RBD datasets showed good correlation with neutralization with *R^2^* up to 0.57 (Fig. 3A). Note, however, that these *R^2^* values cannot be directly compared with *R^2^* values for the convalescent data, because the two sets of analyses are based on distinct datasets. In contrast, the IgM datasets exhibited much poorer correlation than the IgG datasets (Fig. 3A), unlike what we observed for the convalescent data (Fig. 2). Prediction with machine learning algorithms trained on the convalescent data still gave higher correlations, with elastic net yielding *R*^2^ of 0.66 and BRANNs 0.59, than the *R*^2^ values with individual isotype-antigen datasets, although these improvements are smaller than those observed with the convalescent data. These results suggest that the algorithms are effective in predicting neutralization capacity of samples that are substantially different from the training set.

**Figure 3.**
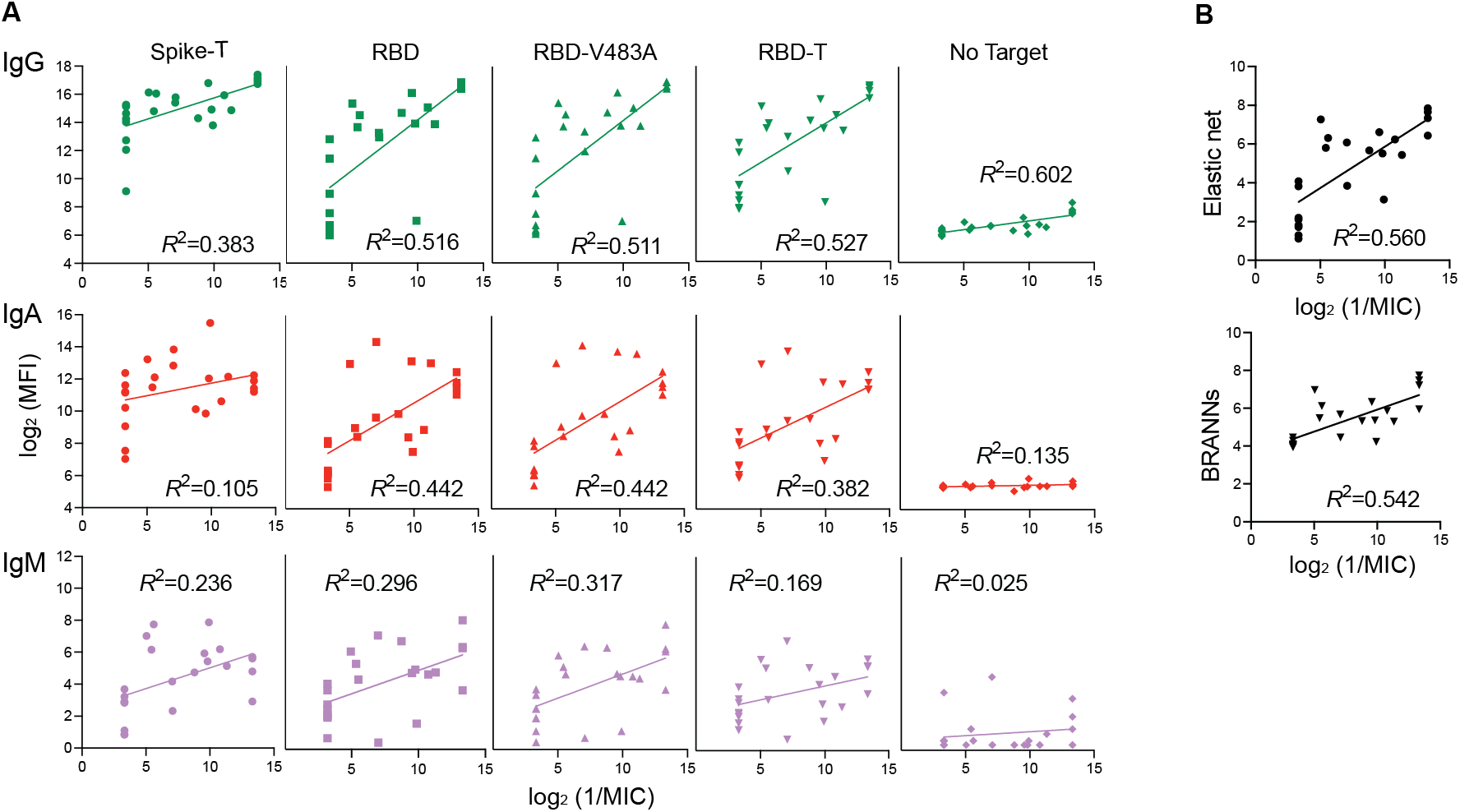
2D-MBBA data for vaccinated lymphoma patients and machine learning predictions of their neutralization capacity. (A) 2D-MBBA data plotted *versus* neutralization titers. Correlation coefficients are also shown. (B) Neutralization titers predicted from machine learning algorithms plotted *versus* experimental titers.

## DISCUSSION

We have developed a high-throughput assay to deeply characterize antibody profiles for SARS-CoV-2 Spike and effective predictive algorithms linking antibody profiles and neutralization capacity. Our assay extends one-dimensional multiplex bead binding assays that have been developed by several groups in binding to multiple antigens is measured for one antibody type at a time (den Hartog et al., 2020; Dobano et al., 2021; Dogan et al., 2021; Garcia-Basteiro et al., 2020; Marien et al., 2021; Pisanic et al., 2020; Weiss et al., 2020; Yates et al., 2021). It is notable that the machine learning algorithms achieved high levels of accuracy from data with a limited set of distinct antigens. Of the final four antigens, RBD and RBD-V483A are similar, and thus we essentially had only three distinct antigens. Still, machine learning algorithms extracted “hidden” information.

The use of a multiple antigen panel is effective in advancing our understanding of the nature of neutralizing antibodies. The effectiveness of the predictive algorithms is in agreement with our current knowledge of the diverse nature of neutralizing antibodies, beyond those targeting the ACE2-interacting surface of the RBD (Amanat et al., 2021; Chi et al., 2020; Liu et al., 2020; Voss et al., 2021). Consistent with this view, the correlation achieved with the predictive algorithms (R^2^ = 0.74) were substantially higher than the results on the same serum sample set for pseudotype virus neutralization assay focusing on the RBD-ACE2 interaction (*R*^2^ = 0.46) (PMID 33692390).

Although the primary goal of this study has been technology development, we have uncovered interesting differences between the convalescent samples and vaccinated lymphoma patient samples. Whereas the ranges of the IgG levels are generally similar between the two sample sets, the vaccinated lymphoma samples had substantially lower IgM levels than the convalescent samples and these IgM datasets showed poor correlation with neutralization. Further studies are needed to pinpoint factors contributing to this difference, such as the underlying disease and treatment and the length of exposure to the infection or vaccination.

There are clear pathways to further improve the 2D-MBBA for SARS-CoV-2 from the version that is described here. New antigens can be added to the panel based on rapidly advancing knowledge of the spike structure and function, e.g., the N-terminal domain and designed mutations that block a particular area of interest, equivalent to the triple mutant antigens. Adding antigens from viral variants of concern (VOCs) would be particularly interesting and probably informative in evaluating the neutralization capacity against such variants. It will be interesting to test how much modification is needed to accurately predict neutralization against VOCs from 2D-MBBA data recorded with antigens containing VOC mutations. Similarly, the assay can be modified to detect different types of antibodies, such as IgG subtypes and Fc receptors (Yates et al., 2021). The potential to accurately predict the neutralization capacity by machine learning of antibody profiling data may contribute to evaluation of vaccines and assessment of high-risk vaccinated populations, particularly in light of recent reports suggesting that the neutralization capacity is a good correlate of protection against SARS-CoV-2 (Earle et al., 2021; Khoury et al., 2021).

The approach employed here can be readily applied to other viruses. Iterative processes of antibody profiling, machine learning prediction of neutralization and updating the antigen panel may be effective in establishing an accurate prediction method for neutralization and in advancing a mechanistic understanding of the nature of viral neutralization and virus-host interactions.

## ACKNOWLEDGMENTS

We would like to thank all the volunteers for participation, Drs. Maria E. Aguero-Rosenfeld and Joan Cangiarella and NYU Langone Hospital Outpatient Laboratory for serum sample collection, and Dr. Meike Dittmann at the NYU Grossman School of Medicine for use of the CX7 CellInsight high-content microscope.

This work was supported in part by the National Institute of Health grants R21 AI158997, R01 CA194864 and R01 CA212608 to S.K., and R21 CA246457 to T.H. K.W.T. was supported by the American Cancer Society fellowship PF-18-180-01-TBE.

## AUTHOR CONTRIBUTIONS

Conceptualization, SK; Methodology, AK, TP, CW, FZL, PA, KAS, HL and SK; Investigation, AK, TP, CW, SAT, LAR, KWT, FZL, PA, ADC, TH, KAS, HL and SK; Resources, JC, CD, MIS and MJM; Writing – original draft, AK, TP, CW and SK; Writing – review and editing, all authors; Supervision, TH, KAS, HL and SK.

## METHODS

### Serum samples

The convalescent serum samples and healthy control samples were previously described (Noval et al., 2021). The vaccinate lymphoma patient samples were described previously (Diefenbach et al., 2021). All patients gave written informed consent and all samples were deidentified for this study under IRB #i20-00595 (SARS-CoV-2 infected), IRB #s18-02037 (healthy pre-SARS-CoV-2 controls), and IRB #S20-02069 (vaccinated lymphoma patients).

### Antigen design, protein expression and purification

The mutations in the RBD were identified from mining the Global Initiative on Sharing All Influenza Data (GISAID) resources. The analysis was based on full genome DNA sequences downloaded on April 13, 2020 (8,086 entries) and May 13, 2020 (24,681 entries). Vector construction has been described previously (Hattori et al., 2021).

The expression vector for the prefusion-stabilized form of the ectodomain of the spike protein (resides 16–1213), SARS-CoV-2 spike RBD (residues 328-531) and SARS-CoV spike RBD (residues 315–517) were described previously (Hattori et al., 2020). The vectors for the mutant RBDs were produced by modifying the vector for wild-type RBD.

All proteins were expressed in Expi293F cells using the recommended transfection kit and the Expi293 Expression Medium (all from ThermoFisher). Briefly, the cells were expanded to 2×10^6^ cells/ml and transfected with 1 ug of DNA per 1 ml of culture volume. The SARS-CoV-2 Spike protein was expressed at 32 °C with 5% CO2 for 4 days. The SARS-COV-2 RBD proteins were expressed at 37 °C with 8% CO2 for 7 days. The culture supernatants were harvested by centrifugation, supplemented with protease inhibitors and clarified by further centrifugation at 8000 rpm for 20 mins as well as filtration through a 0.22 μm filter. The supernatants were dialyzed into 20 mM sodium phosphate pH 7.4 with 500 mM sodium chloride and purified using HisTrap Excel affinity column (GE). After HisTrap Excel purification, the proteins were biotinylated using an in-house purified BirA enzyme. To remove residual BirA enzyme, the RBD proteins were subjected to an additional HisTrap Excel purification step and dialyzed into 1x PBS. The residual BirA enzyme was removed from the spike protein samples by size exclusion chromatography using a Superdex200 Increase column in 20 mM sodium phosphate pH 6.3 with 500 mM sodium choloride. All proteins were snap frozen and stored at −80 °C.

### Multiplex Bead Binding Assay

QBeads DevScreen: SAv (Streptavidin) (Sartorius 90792) were used to immobilize biotinylated Spike, RBD domains and the associated mutants. The Qbeads kit is composed of five polystyrene bead types that differ in the amount of internal fluorescence they contain but otherwise identical. Bead preparation started with appropriate volumes of stock beads that give 500 Qbeads of each type per reaction. For each bead type, the beads were washed twice with 1x PBS containing 0.5% (w/v) BSA (PBS-BSA) and resuspended in the starting bead volume by following the vendor’s instructions. A biotinylated antigen protein at 25 nM in the volume equal to that of the suspended beads was added to the washed beads and incubated for 30 min at 4°C, then 10 μM biotin in PBS-BSA in the volume equal to that of the antigen solution was added and incubated for 10 min at 4°C to block unoccupied biotin-binding sites of streptavidin. The beads were then washed twice by spinning down at 8000 x g for 5 minutes and resuspending in PBS-BSA containing 10 μM biotin.

Different bead types loaded with antigens were then combined and diluted in enough buffer to allow the distribution of 5 μl of bead mixture per well of a 96 well HTS filter plate (Thermo Fisher, catalog number MSHVN4550) using a MANTIS dispensing robot (Formulatrix). The beads were dispensed into HTS 96 well filter plates that have been pre-washed and preloaded with 7.5 μl of filtered 1% non-fat dry milk solution in PBS containing 0.1% Tween20 (PBST).

Serum samples were heat-inactivated by incubating at 58 °C for 1 hr, centrifuged to eliminate debris, diluted five-fold in PBS, aliquoted and flash-frozen in liquid nitrogen and kept at −80°C until the day of the assay. The frozen samples were thawed, diluted 5:79 in PBST containing 1% milk. Then 12.5 μl of serum were added to each well. For reference standards, a commercially available anti-COVID-19 and SARS-CoV S glycoprotein antibody clone CR3022 in the IgG, IgA and IgM formats (Absolute Antibody, Human IgG1, Kappa, catalog number Ab01680-10.0, Human IgA, Kappa, catalog number Ab01680-16.0, Human IgM, Kappa, catalog number Ab01680-15.0) were included in triplicates in each measurement. The plates were incubated at room temperature for 30 min, washed three times with 1x PBST containing 0.1% BSA using a vacuum chamber, and stained with 50 μl of secondary antibody mixture for 30 minutes. Anti-human IgG FITC (Jackson 109-545-098 diluted 1:800), anti-human IgA PE (Jackson 109-115-011 diluted 1:100) and anti-human IgM DyLight405 (Jackson 709-475-073 diluted 1:200) were used as secondary antibodies. After the secondary antibody incubation, the plates were washed twice with PBST containing 0.1% BSA and resuspended in 80 μl wash buffer, measured using a Yeti ZE5 Cell Analyzer (Bio-Rad). Data were analyzed using FlowJo (BD, version 10.7.1).

In order to standardize MBBA data across different measurements, they were referenced to the MFI values of the control antibodies, CR3022, as described above. All MFI values in each measurement set, i.e., on a single 96-well plate, were scaled so that the mean MFI values 6 nM CR3022-IgG1, 2 nM CR3022-IgA and 4 nM CR3022-IgM are 37653, 17152 and 1144, respectively, for Spike and that the mean MFI values of these antibodies for RBD were 106436, 37076 and 3576, respectively.

### Neutralization assay

Neutralization values for the convalescent serum samples were taken from a previous publication (Noval et al., 2021). For neutralization of cancer patient serum, 20,000 Vero E6 (ATCC CRL-1586) cells/well were seeded in 96 well plates the day before infection. Patient serum was 2-fold serially diluted (ranging from 1:20 to 1:10,240) in DMEM supplemented with 1% nonessential amino acids, 10 mM HEPES, and 2% fetal bovine serum. Each serum dilution was mixed 1:1 (vol/vol) with 5000 PFU of icSARS-CoV-2-mNG (a gift from the UTMB World Reference Center for Emerging Viruses and Arboviruses (PDMI: 32289263)) and incubated at 37°C for 1 hr. The virus:serum mix was then added to the Vero E6 cell containing plates and incubated at 37°C for 24 hrs. After incubation, cells were fixed with 10% formalin, stained with DAPI, and virus positive cells were quantified using a CellInsight CX7 High-content microscope using a cut-off of three standard deviations from negative to be scored as an infected cell. All neutralization assays were performed in duplicate. Samples were compared to an untreated virus control.

### Machine learning prediction of neutralization capacity

Neutralization and assay parameters were first log2-transformed. We employed multiple well-known machine learning algorithms: stepwise linear regression, elastic net regression, multivariate adaptive regression splines, classification and regression tree, and Bayesian regularized neural networks, to establish an accurate predictive model for neutralization capacity from serology data. Repeated 10-fold CV was employed to evaluate the predictive performance of these algorithms. Specifically, the sample set was randomly split into a training set with 90% observations and a test set with 10% observations. The training set was employed to build the final predictive model and the test set was employed to calculate the predictive metrics RMSE, *R^2^*, and MAE with the trained model, for each algorithm. For the algorithms consisting of tuning parameters, 10-fold CV was employed to identify the best tuning parameters with the lowest RMSE. The above procedure was replicated 100 times to robustly determine the best predictive model for neutralization (less dependent on the split in training and test sets).

In terms of the three evaluation metrics above, BRANNs and elastic net regression have the best performance on predicting neutralization among all predictive algorithms. We further identify crucial serology parameters that have important roles in predicting neutralization capacity in these two methods, respectively. Specifically, for BRANNs, we employ the connection weights method (Olden et al., 2004) to determine the variable importance, which is a more flexible approach to evaluate variable importance in neural networks. This method calculates importance as the sum of the product of the input-hidden and hidden-output connection weights across all hidden neurons, which can be capable of evaluating neural networks with multiple hidden layers and response variables. On the other hand, we employed the regression coefficients to determine the variable importance for elastic net regression. Both methods maintain in terms of both magnitude and sign. The sign indicates the direction of correlation (negative or positive), and the absolute value indicates the magnitude of importance. The importance rank (relative importance) is calculated according to the absolute connection weights in BARNNs or the absolute estimated coefficients in elastic net regression.

## SUPPLEMENTARY INFORMATION

Figure S1. Preparation of antigen samples.

Figure S2. Validation of 2D-MBBA method.

Figure S3. The receiver operating characteristic (ROC) curves of individual datasets.

Figure S4. Correlation among 2D-MBBA datasets and their rankings in neutralization prediction.

## REFERENCES

Agha, M., Blake, M., Chilleo, C., Wells, A., and Haidar, G. (2021). Suboptimal response to COVID-19 mRNA vaccines in hematologic malignancies patients. medRxiv, 10.1101/2021.1104.1106.21254949.

Amanat, F., Stadlbauer, D., Strohmeier, S., Nguyen, T.H.O., Chromikova, V., McMahon, M., Jiang, K., Arunkumar, G.A., Jurczyszak, D., Polanco, J., et al. (2020). A serological assay to detect SARS-CoV-2 seroconversion in humans. Nat Med 26, 1033–1036.

Amanat, F., Thapa, M., Lei, T., Ahmed, S.M.S., Adelsberg, D.C., Carreno, J.M., Strohmeier, S., Schmitz, A.J., Zafar, S., Zhou, J.Q., et al. (2021). SARS-CoV-2 mRNA vaccination induces functionally diverse antibodies to NTD, RBD, and S2. Cell 184, 3936–3948.

Baden, L.R., El Sahly, H.M., Essink, B., Kotloff, K., Frey, S., Novak, R., Diemert, D., Spector, S.A., Rouphael, N., Creech, C.B., et al. (2021). Efficacy and Safety of the mRNA-1273 SARS-CoV-2 Vaccine. N Engl J Med 384, 403–416.

Bewley, K.R., Coombes, N.S., Gagnon, L., McInroy, L., Baker, N., Shaik, I., St-Jean, J.R., St-Amant, N., Buttigieg, K.R., Humphries, H.E., et al. (2021). Quantification of SARS-CoV-2 neutralizing antibody by wild-type plaque reduction neutralization, microneutralization and pseudotyped virus neutralization assays. Nat Protoc 16, 3114–3140.

Bojkova, D., Klann, K., Koch, B., Widera, M., Krause, D., Ciesek, S., Cinatl, J., and Munch, C. (2020). Proteomics of SARS-CoV-2-infected host cells reveals therapy targets. Nature 583, 469–472.

Boyarsky, B.J., Werbel, W.A., Avery, R.K., Tobian, A.A.R., Massie, A.B., Segev, D.L., and Garonzik-Wang, J.M. (2021). Immunogenicity of a Single Dose of SARS-CoV-2 Messenger RNA Vaccine in Solid Organ Transplant Recipients. JAMA 325, 1784–1786.

Brouwer, P.J.M., Caniels, T.G., van der Straten, K., Snitselaar, J.L., Aldon, Y., Bangaru, S., Torres, J.L., Okba, N.M.A., Claireaux, M., Kerster, G., et al. (2020). Potent neutralizing antibodies from COVID-19 patients define multiple targets of vulnerability. Science 369, 643–650.

Burden, F., and Winkler, D. (2008). Bayesian regularization of neural networks. Methods Mol Biol 458, 25–44.

Chi, X., Yan, R., Zhang, J., Zhang, G., Zhang, Y., Hao, M., Zhang, Z., Fan, P., Dong, Y., Yang, Y., et al. (2020). A neutralizing human antibody binds to the N-terminal domain of the Spike protein of SARS-CoV-2. Science 369, 650–655.

Deepak, P., Kim, W., Paley, M.A., Yang, M., Carvidi, A.B., El-Qunni, A.A., Haile, A., Huang, K., Kinnett, B., Liebeskind, M.J., et al. (2021). Glucocorticoids and B Cell Depleting Agents Substantially Impair Immunogenicity of mRNA Vaccines to SARS-CoV-2. medRxiv, 10.1101/2021.1104.1105.21254656.

den Hartog, G., Schepp, R.M., Kuijer, M., GeurtsvanKessel, C., van Beek, J., Rots, N., Koopmans, M.P.G., van der Klis, F.R.M., and van Binnendijk, R.S. (2020). SARS-CoV-2-Specific Antibody Detection for Seroepidemiology: A Multiplex Analysis Approach Accounting for Accurate Seroprevalence. J Infect Dis 222, 1452–1461.

Diefenbach, C., Caro, J., Koide, A., Grossbard, M., Goldberg, J.D., Raphael, B., Hymes, K., Moskovits, T., Kreditor, M., Kaminetzky, D., et al. (2021). Impaired Humoral Immunity to SARS-CoV-2 Vaccination in Non-Hodgkin Lymphoma and CLL Patients. medRxiv, 10.1101/2021.1106.1102.21257804.

Dobano, C., Vidal, M., Santano, R., Jimenez, A., Chi, J., Barrios, D., Ruiz-Olalla, G., Rodrigo Melero, N., Carolis, C., Parras, D., et al. (2021). Highly Sensitive and Specific Multiplex Antibody Assays To Quantify Immunoglobulins M, A, and G against SARS-CoV-2 Antigens. J Clin Microbiol 59, e01731–01720.

Dogan, M., Kozhaya, L., Placek, L., Gunter, C., Yigit, M., Hardy, R., Plassmeyer, M., Coatney, P., Lillard, K., Bukhari, Z., et al. (2021). SARS-CoV-2 specific antibody and neutralization assays reveal the wide range of the humoral immune response to virus. Commun Biol 4, 129.

Earle, K.A., Ambrosino, D.M., Fiore-Gartland, A., Goldblatt, D., Gilbert, P.B., Siber, G.R., Dull, P., and Plotkin, S.A. (2021). Evidence for antibody as a protective correlate for COVID-19 vaccines. Vaccine 39, 4423–4428.

Friedman, J.H. (1991). Multivariate adaptive regression splines. Ann Statist 19, 1–67.

Garcia-Basteiro, A.L., Moncunill, G., Tortajada, M., Vidal, M., Guinovart, C., Jimenez, A., Santano, R., Sanz, S., Mendez, S., Llupia, A., et al. (2020). Seroprevalence of antibodies against SARS-CoV-2 among health care workers in a large Spanish reference hospital. Nat Commun 11, 3500.

Hansen, J., Baum, A., Pascal, K.E., Russo, V., Giordano, S., Wloga, E., Fulton, B.O., Yan, Y., Koon, K., Patel, K., et al. (2020). Studies in humanized mice and convalescent humans yield a SARS-CoV-2 antibody cocktail. Science 369, 1010–1014.

Hattori, T., Koide, A., Panchenko, T., Romero, L.A., Teng, K.W., Corrado, A.D., and Koide, S. (2020). Multiplex bead binding assays using off-the-shelf components and common flow cytometers. J Immunol Methods 490, 112952.

Hattori, T., Koide, A., Panchenko, T., Romero, L.A., Teng, K.W., Tada, T., Landau, N.R., and Koide, S. (2021). The ACE2-binding interface of SARS-CoV-2 Spike inherently deflects immune recognition. J Mol Biol 433, 166748.

Herishanu, Y., Avivi, I., Aharon, A., Shefer, G., Levi, S., Bronstein, Y., Morales, M., Ziv, T., Shorer Arbel, Y., Scarfo, L., et al. (2021). Efficacy of the BNT162b2 mRNA COVID-19 vaccine in patients with chronic lymphocytic leukemia. Blood 137, 3165–3173.

Hoffmann, M., Kleine-Weber, H., Schroeder, S., Kruger, N., Herrler, T., Erichsen, S., Schiergens, T.S., Herrler, G., Wu, N.H., Nitsche, A., et al. (2020). SARS-CoV-2 Cell Entry Depends on ACE2 and TMPRSS2 and Is Blocked by a Clinically Proven Protease Inhibitor. Cell 181, 271–280 e278.

Jackson, L.A., Anderson, E.J., Rouphael, N.G., Roberts, P.C., Makhene, M., Coler, R.N., McCullough, M.P., Chappell, J.D., Denison, M.R., Stevens, L.J., et al. (2020). An mRNA Vaccine against SARS-CoV-2 - Preliminary Report. N Engl J Med 383, 1920–1931.

Ju, B., Zhang, Q., Ge, J., Wang, R., Sun, J., Ge, X., Yu, J., Shan, S., Zhou, B., Song, S., et al. (2020). Human neutralizing antibodies elicited by SARS-CoV-2 infection. Nature 584, 115–119.

Khoury, D.S., Cromer, D., Reynaldi, A., Schlub, T.E., Wheatley, A.K., Juno, J.A., Subbarao, K., Kent, S.J., Triccas, J.A., and Davenport, M.P. (2021). Neutralizing antibody levels are highly predictive of immune protection from symptomatic SARS-CoV-2 infection. Nat Med 27, 1205–1211.

Lan, J., Ge, J., Yu, J., Shan, S., Zhou, H., Fan, S., Zhang, Q., Shi, X., Wang, Q., Zhang, L., et al. (2020). Structure of the SARS-CoV-2 spike receptor-binding domain bound to the ACE2 receptor. Nature 581, 215–220.

Lewis, R.J. (2000). An introduction to classification and regression tree (CART) analysis. Annu Meeting Soc Acad Mergency Med 14.

Liu, L., Wang, P., Nair, M.S., Yu, J., Rapp, M., Wang, Q., Luo, Y., Chan, J.F., Sahi, V., Figueroa, A., et al. (2020). Potent neutralizing antibodies against multiple epitopes on SARS-CoV-2 spike. Nature 584, 450–456.

Long, Q.X., Liu, B.Z., Deng, H.J., Wu, G.C., Deng, K., Chen, Y.K., Liao, P., Qiu, J.F., Lin, Y., Cai, X.F., et al. (2020). Antibody responses to SARS-CoV-2 in patients with COVID-19. Nat Med 26, 845–848.

Marien, J., Ceulemans, A., Michiels, J., Heyndrickx, L., Kerkhof, K., Foque, N., Widdowson, M.A., Mortgat, L., Duysburgh, E., Desombere, I., et al. (2021). Evaluating SARS-CoV-2 spike and nucleocapsid proteins as targets for antibody detection in severe and mild COVID-19 cases using a Luminex bead-based assay. J Virol Methods 288, 114025.

Miller, A. (2002). Subset selection in regression (CRC Press).

Muruato, A.E., Fontes-Garfias, C.R., Ren, P., Garcia-Blanco, M.A., Menachery, V.D., Xie, X., and Shi, P.Y. (2020). A high-throughput neutralizing antibody assay for COVID-19 diagnosis and vaccine evaluation. Nat Commun 11, 4059.

Nie, J., Li, Q., Wu, J., Zhao, C., Hao, H., Liu, H., Zhang, L., Nie, L., Qin, H., Wang, M., et al. (2020). Quantification of SARS-CoV-2 neutralizing antibody by a pseudotyped virus-based assay. Nat Protoc 15, 3699–3715.

Noval, M.G., Kaczmarek, M.E., Koide, A., Rodriguez-Rodriguez, B.A., Louie, P., Tada, T., Hattori, T., Panchenko, T., Romero, L.A., Teng, K.W., et al. (2021). Antibody isotype diversity against SARS-CoV-2 is associated with differential serum neutralization capacities. Sci Rep 11, 5538.

Okba, N.M.A., Muller, M.A., Li, W., Wang, C., GeurtsvanKessel, C.H., Corman, V.M., Lamers, M.M., Sikkema, R.S., de Bruin, E., Chandler, F.D., et al. (2020). Severe Acute Respiratory Syndrome Coronavirus 2-Specific Antibody Responses in Coronavirus Disease 2019 Patients. Emerg Infect Dis 26, 1478–1488.

Olden, J.D., Joy, M.K., and Death, R.G. (2004). An accurate comparison of methods for quantifying variable importance in artificial neural networks using simulated data. Ecol Modelling 178, 389–397.

Pisanic, N., Randad, P.R., Kruczynski, K., Manabe, Y.C., Thomas, D.L., Pekosz, A., Klein, S.L., Betenbaugh, M.J., Clarke, W.A., Laeyendecker, O., et al. (2020). COVID-19 Serology at Population Scale: SARS-CoV-2-Specific Antibody Responses in Saliva. J Clin Microbiol 59, e02204–02220.

Polack, F.P., Thomas, S.J., Kitchin, N., Absalon, J., Gurtman, A., Lockhart, S., Perez, J.L., Perez Marc, G., Moreira, E.D., Zerbini, C., et al. (2020). Safety and Efficacy of the BNT162b2 mRNA Covid-19 Vaccine. N Engl J Med 383, 2603–2615.

Riepler, L., Rossler, A., Falch, A., Volland, A., Borena, W., von Laer, D., and Kimpel, J. (2020). Comparison of Four SARS-CoV-2 Neutralization Assays. Vaccines (Basel) 9, 13.

Robbiani, D.F., Gaebler, C., Muecksch, F., Lorenzi, J.C.C., Wang, Z., Cho, A., Agudelo, M., Barnes, C.O., Gazumyan, A., Finkin, S., et al. (2020). Convergent antibody responses to SARS-CoV-2 in convalescent individuals. Nature 584, 437–442.

Shi, R., Shan, C., Duan, X., Chen, Z., Liu, P., Song, J., Song, T., Bi, X., Han, C., Wu, L., et al. (2020). A human neutralizing antibody targets the receptor-binding site of SARS-CoV-2. Nature 584, 120–124.

Tan, C.W., Chia, W.N., Qin, X., Liu, P., Chen, M.I., Tiu, C., Hu, Z., Chen, V.C., Young, B.E., Sia, W.R., et al. (2020). A SARS-CoV-2 surrogate virus neutralization test based on antibody-mediated blockage of ACE2-spike protein-protein interaction. Nat Biotechnol 38, 1073–1078.

Voss, W.N., Hou, Y.J., Johnson, N.V., Delidakis, G., Kim, J.E., Javanmardi, K., Horton, A.P., Bartzoka, F., Paresi, C.J., Tanno, Y., et al. (2021). Prevalent, protective, and convergent IgG recognition of SARS-CoV-2 non-RBD spike epitopes. Science 372, 1108–1112.

Voysey, M., Clemens, S.A.C., Madhi, S.A., Weckx, L.Y., Folegatti, P.M., Aley, P.K., Angus, B., Baillie, V.L., Barnabas, S.L., Bhorat, Q.E., et al. (2021). Safety and efficacy of the ChAdOx1 nCoV-19 vaccine (AZD1222) against SARS-CoV-2: an interim analysis of four randomised controlled trials in Brazil, South Africa, and the UK. Lancet 397, 99–111.

Walls, A.C., Park, Y.J., Tortorici, M.A., Wall, A., McGuire, A.T., and Veesler, D. (2020). Structure, Function, and Antigenicity of the SARS-CoV-2 Spike Glycoprotein. Cell 181, 281–292 e286.

Walsh, E.E., Frenck, R.W., Jr., Falsey, A.R., Kitchin, N., Absalon, J., Gurtman, A., Lockhart, S., Neuzil, K., Mulligan, M.J., Bailey, R., et al. (2020). Safety and Immunogenicity of Two RNA-Based Covid-19 Vaccine Candidates. N Engl J Med 383, 2439–2450.

Wec, A.Z., Wrapp, D., Herbert, A.S., Maurer, D.P., Haslwanter, D., Sakharkar, M., Jangra, R.K., Dieterle, M.E., Lilov, A., Huang, D., et al. (2020). Broad neutralization of SARS-related viruses by human monoclonal antibodies. Science 369, 731–736.

Weiss, S., Klingler, J., Hioe, C., Amanat, F., Baine, I., Arinsburg, S., Kojic, E.M., Stoever, J., Liu, S.T.H., Jurczyszak, D., et al. (2020). A High-Throughput Assay for Circulating Antibodies Directed Against the S Protein of Severe Acute Respiratory Syndrome Coronavirus 2. J Infect Dis 222, 1629–1634.

Woof, J.M., and Russell, M.W. (2011). Structure and function relationships in IgA. Mucosal Immunol 4, 590–597.

Wu, Y., Wang, F., Shen, C., Peng, W., Li, D., Zhao, C., Li, Z., Li, S., Bi, Y., Yang, Y., et al. (2020). A noncompeting pair of human neutralizing antibodies block COVID-19 virus binding to its receptor ACE2. Science 368, 1274–1278.

Xie, X., Muruato, A., Lokugamage, K.G., Narayanan, K., Zhang, X., Zou, J., Liu, J., Schindewolf, C., Bopp, N.E., Aguilar, P.V., et al. (2020). An Infectious cDNA Clone of SARS-CoV-2. Cell Host Microbe 27, 841–848 e843.

Yates, J.L., Ehrbar, D.J., Hunt, D.T., Girardin, R.C., Dupuis, A.P., 2nd, Payne, A.F., Sowizral, M., Varney, S., Kulas, K.E., Demarest, V.L., et al. (2021). Serological Analysis Reveals an Imbalanced IgG Subclass Composition Associated with COVID-19 Disease Severity. Cell Rep Med, 100329.

Zou, H., and Hastie, T. (2005). Regularization and variable selection via the elastic net. J R Statist Soc B 67, 301–320.

